# Remodeling tumor microenvironment by liposomal co-delivery of DMXAA and simvastatin inhibits malignant melanoma progression

**DOI:** 10.1101/2021.04.26.441259

**Authors:** Valentin-Florian Rauca, Laura Patras, Lavinia Luput, Emilia Licarete, Vlad-Alexandru Toma, Alina Porfire, Augustin-Catalin Mot, Elena Rakosy-Tican, Alina Sesarman, Manuela Banciu

## Abstract

Anti-angiogenic therapies for melanoma have not yet been translated into meaningful clinical benefit for patients, due to development of drug-induced resistance in cancer cells, mainly caused by hypoxia-inducible factor 1α (HIF-1α) overexpression and enhanced oxidative stress mediated by tumor-associated macrophages (TAMs). Our previous study demonstrated synergistic antitumor actions of simvastatin (SIM) and 5,6-dimethylxanthenone-4-acetic acid (DMXAA) on an *in vitro* melanoma model *via* suppression of the aggressive phenotype of melanoma cells and inhibition of TAMs-mediated angiogenesis. Therefore, we took the advantage of long circulating liposomes (LCL) superior tumor targeting capacity to efficiently deliver SIM and DMXAA to B16.F10 melanoma *in vivo*, with the final aim of improving the outcome of the anti-angiogenic therapy. Thus, we assessed the effects of this novel combined tumor-targeted treatment on *s*.*c*. B16.F10 murine melanoma growth and on the production of critical markers involved in tumor development and progression. Our results showed that the combined liposomal therapy inhibited almost totally the growth of melanoma tumors, due to the enhancement of anti-angiogenic effects of LCL-DMXAA by LCL-SIM and induction of a pro-apoptotic state in the tumor microenvironment (TME). These effects were favoured by the partial re-education of TAMs towards a M1 phenotype and maintained via suppression of major invasion and metastasis promoters (HIF-1α, pAP-1 c-Jun, and MMPs). Thus, this novel therapy holds the potential to remodel the tumor microenvironment, by suppressing its most important malignant biological capabilities.

**Highlights:**

- Novel combined liposomal delivery of SIM and DMXAA inhibits melanoma tumor growth
- LCL-SIM augments the anti-angiogenic effects of LCL-DMXAA
- Combined liposomal therapy inhibits HIF-1α/VEGF axis, induces apoptosis and TAM re-education in tumors
- This novel therapy suppresses the most important malignant capabilities of melanoma

## Introduction

Melanoma cells are established providers of essential growth factors to trigger tumor angiogenesis, such as VEGF and bFGF that further support tumor development and metastasis [1]. Therefore, targeting tumor vasculature with anti-angiogenic drugs such as vascular disrupting agents (VDA) seemed like a promising approach in the treatment of solid tumors, albeit drug resistance associated with anti-angiogenic therapy was reported in most of the cases [2,3]. Especially after VDA treatment, a remaining viable tumor rim, characterized by intratumor overexpression of hypoxia-inducible factor 1α (HIF-1α) and enhanced oxidative stress mediated by tumor-associated macrophages (TAMs), is responsible for selecting aggressive tumor cell phenotypes ready to escape oxygen and nutrient deprivation and thus, accelerating the undesired outcome of the disease [3–5]. Moreover, our previous findings have shown that simvastatin (SIM)-a lipophilic statin incorporated in long-circulating liposomes (LCL-SIM) could counteract both causes of cancer resistance to anti-angiogenic treatments as LCL-SIM inhibited B16.F10 murine melanoma growth *in vivo via* suppression of TAMs-mediated oxidative stress and HIF-1α levels in melanoma cells [6]. Thus, the involvement of intratumor macrophages in tumor cell resistance to apoptosis and chemotherapy might be exploited for future TAMs-targeted therapies that can counteract negative outcomes of the anti-angiogenic treatments [7]. Furthermore, in another recent study when we administered SIM in combination with a VDA, 5,6-dimethylxanthenone-4-acetic acid (DMXAA), the aggressiveness of melanoma cells was suppressed due to the synergistic action on cancer cell proliferation as well as inhibition of protumor function of TAMs *in vitro* [8]. In tight connection with these data, it has been shown recently that the combination between an anti-angiogenic agent and an antioxidant modulator could counteract the effect of HIF on cancer cell metabolism [9]. Thus, targeting the mediators of communication between cancer cell and cells residing the TME may successfully complement other treatment alternatives [10].

In the present study, we aimed to improve the outcome of anti-angiogenic therapy for B16.F10 melanoma *in vivo* by using a novel tumor-targeted approach based on co-delivery of liposomal DMXAA together with liposomal SIM. To our knowledge, this therapeutic approach has never been described before. Thus, we evaluated the effects of this combined tumor-targeted treatment on *s*.*c*. B16.F10 murine melanoma growth, with regard to the levels of specific markers involved in angiogenesis, inflammation, oxidative stress, apoptosis and invasion and metastasis. Our results showed that this novel targeted therapy holds the potential to remodel the TME, by suppressing its most important malignant biological capabilities.

## Materials and methods

### Preparation and physicochemical characterization of liposomal formulations

DPPC and PEG-2000-DSPE were acquired from Lipoid GmbH (Ludwigshafen, GER), CHL and SIM from Sigma-Aldrich Chemie GmbH (Munich, GER) and DMXAA was purchased from Selleck Chemicals LLC (Houston, TX). The molar ratio of compounds used for LCL-SIM preparation was 17:1.01:1:1.209 (DPPC:PEG-2000-DSPE:CHL:SIM), according to our previous published protocols [6]. The molar ratio of compounds used for the preparation of the novel DMXAA liposomal formulation was 1.85:0.7:0.3:0.15 (DPPC:CHL:DMXAA:PEG-2000-DSPE) and was based on previous studies regarding nanoformulations that encapsulated small molecule therapeutic agents [11,12]. Lipid film hydration method followed by multiple extrusion steps was used to prepare nanoliposomes as described previously [6]. Each LCL formulation was characterized as size, polydispersity index, zeta potential, the concentration of the active drug, and their entrapment efficiencies.

### Cell type and murine tumor model

B16.F10 murine melanoma cells (ATCC, CRL-6475) were cultured in DMEM according to our previously described methods [8]. Syngeneic male C57BL/6 mice 6 to 8-week-old (Cantacuzino Institute, Bucharest, RO) kept under standard laboratory conditions were inoculated with 1 × 10^6^ B16.F10 cells *s*.*c*. in the right flank. Tumors were measured daily and tumor volume was calculated as described before [13]. Treatments started at day 11 after cell inoculation when tumors were about 100 mm^3^. Animal experiments were performed according to the EU Directive 2010/63/EU and to the national regulations and were approved by the Babes-Bolyai University Ethics Committee (Cluj-Napoca, Romania; 4335/19.03.2018).

### Effects of different treatments on tumor growth

The effects of liposome-encapsulated agents SIM and DMXAA on tumor growth were compared to the effects of free active agents on B16.F10 murine melanoma-bearing mice. LCL-SIM (5 mg/kg) and LCL-DMXAA (14 mg/kg) were administered as monotherapies or combined, in the caudal vein of C57BL/6 mice, on days 11 and 14 after tumor cell inoculation. Each experimental group consisted of 5 animals and the control group was treated with LCL (*i*.*e*. devoid of drug). Tumor size and body weight were monitored on a daily basis during treatment. Tumor volume was assessed using the formula V = 0.52*a*^*2*^*b*, where *a* is the smallest and *b* is the largest superficial diameter of the tumor. Mice from all experimental groups were sacrificed on day 15 and tumors were collected and stored in liquid nitrogen until analysis.

### Angiogenic/ inflammatory protein array analysis

To determine the effect of single and co-administered liposomal therapies on the expression levels of angiogenic/inflammatory proteins in whole tumor lysates, a screening for 24 proteins involved in these major protumoral processes was performed, using the RayBio^®^ Mouse Angiogenic protein Antibody Array membranes 1.1 (RayBiotech Inc., Norcross, GA, USA) as previously detailed [8,12].

### Histology and immunohistochemistry analysis

For the evaluation of histological and immunohistochemical features, the tumors were processed as previously described [13]. After paraffin embedding, sections were cut at 5 µm and mounted on positively charged glass slides. The primary antibody-rabbit polyclonal IgG anti-mouse CD31 (ab124432, Abcam, Cambridge, UK) was diluted 1000-fold. The slides were examined by light microscopy and the positive reaction (area of the brown staining) was evaluated in six different microscope fields. We used the following scoring system to evaluate the area percentage (%) of CD-31 positive immunoreaction: 0.5 -5-20%; 1 -20-40%; 2 -40-60%; 3 -60-80%; 4 -80-100%.

### Western Blot quantification of tumor tissue proteins

Frozen tumors from each experimental groups were pooled to obtain tumor tissue lysates [14]. 5-10 μg proteins from each lysate were separated by SDS-PAGE onto a 10% polyacrylamide gel and immunobloted against the following primary antibodies, diluted 500-fold: mouse monoclonal IgG anti-mouse Bcl-xL (sc-8392, Santa Cruz Biotechnology, Texas, USA), rabbit polyclonal IgG anti-mouse Bax (2772S, Cell Signaling Technology, Inc, Danvers, USA), rabbit monoclonal IgG anti-mouse HIF-1α (ab179483, Abcam, Cambridge, UK), and rabbit polyclonal IgG anti-mouse pAP1-c-Jun (sc-7981-R, Santa Cruz Biotechnology, Texas, USA). β-actin which was used as loading control was detected using a rabbit polyclonal IgG against mouse β-actin (A2103, Merck, Darmstadt, GER) diluted 1000-fold. For detection of the bound antibodies, goat anti-rabbit IgG HRP-labeled (sc-2004, Santa Cruz Biotechnology, Texas, USA) and goat anti-mouse IgG HRP-labeled antibodies (sc-2005, Santa Cruz Biotechnology, Texas, USA) diluted 4000-fold, were used. Protein detection and quantification were performed as previously reported [8].

### Evaluation of oxidative stress parameters

The levels of lipid peroxidation marker MDA were measured by HPLC [15]. Data were normalized to the protein concentration in tumor lysates and expressed as nmoles MDA/mg protein. Intratumor activity of catalase was assessed using the method described by Aebi [16] and expressed as units of catalytic activity/mg protein. The evaluation of total antioxidant capacity (TAC) of the TME was performed according to the method described by Erel [17] and expressed as μmoles Trolox/mg protein. For these assays, each sample was determined in duplicate.

### Gelatin zymography analysis of MMP-2 and MMP-9 activity

Electrophoretic gels containing 0.1% gelatin and 7.5% acrylamide were used to fractionate 30 μg proteins from tumor lysates, under denaturating but non-reducing conditions. Determination of the gelatinolytic activity of MMP-2 and MMP-9 in tumor lysates followed previously published protocols [18].

### RT-qPCR determination of Arg-1 and iNOS mARN expression

Total RNA was isolated from frozen tumors using an RNA kit (peqGOLD Total RNA Kit, PeqLab, Erlangen, DE). To avoid potential DNA contamination, 2 μg of total RNA were digested with 2U of RNase free DNase (Thermo Scientific, MA, USA) for 30 min at 37° C, followed by addition of EDTA and incubation at 65°C for 10 min. From the resulting DNA-free RNA, 1μg was reverse-transcribed into cDNA using Verso cDNA kit (ThermoScientific, MA, USA), while identical samples from each experimental group processed in the absence of reverse transcriptase served as DNA contamination controls, as previously described [8]. Reverse transcription products (1μl) were added to a 25-μl reaction mix containing 1×Maxima SYBR Green qPCR Master Mix (Thermo Scientific, MA, USA) and 0.3 μM of each primer. Real-time PCR reactions were performed under the following cycling parameters: pre-incubation at 95° C for 10 min, cycling: 95° C for 15 s, 60° C for 30 s, and then 72° C for 30 s. Melting curves were generated to check for the primers specificity. The primers used for gene amplification are presented in **Supplementary Table 1**. Comparative Ct method (ΔΔCt) was used to calculate gene expression by relative quantitation. Gene expression was reported as fold change (2^-ΔΔCt^), relative to mARN expression in Control tumors, used as calibrator. Mouse β-actin mRNA was used as reference gene expression.

### Statistical analysis

Data from different experiments were expressed as mean ± standard deviation (SD). All statistical analyses were performed by using GraphPad Prism version 6 for Windows (GraphPad Software, San Diego, CA, USA). The overall effects of different treatments on tumor growth, on intratumor levels of anti-apoptotic proteins, key invasion and metastasis promoters and oxidative stress markers were analyzed by one-way ANOVA with Bonferroni correction for multiple comparisons. For the estimation of the treatments actions on angiogenic and inflammatory protein production, 2-way ANOVA with Bonferroni correction for multiple comparisons was used. The scores for immunoreaction intensities of tumor sections from different experimental groups were analyzed by using rank-based nonparametric Kruskall-Wallis test with Dunn’s test for multiple comparisons. A *P* value of < 0.05 was considered significant.

## Results

### Characterization of liposomal drug formulations

As shown in **Supplementary Table 2**, LCL-SIM and the novel LCL-DMXAA formulation were characterized regarding particle size distribution, polydispersity index, zeta potential, the concentration of the active drug and encapsulation efficiency. Importantly, mean particle size of the liposomes was found to be around 110-135 nm (below the cutoff limits of the pores of tumor endothelia which are 200-800 nm) [19] with a narrow size distribution (polydispersity index lower than 0.1, **Supplementary Table 2**). Thus, given this attributes the liposomal formulation with SIM and DMXAA might have the ability to substantially extravasate and accumulate in tumors due to the enhanced permeability of tumor vasculature (referred to as the EPR “enhanced permeability and retention” effect), as compared to healthy endothelium [6,11]. Notably, the encapsulation efficiencies values are very high for a hydrophobic drug such as SIM (over 80% for LCL-SIM) and for a hydrophilic drug such as DMXAA (about 40% for LCL-DMXAA), suggesting potential for future technological transfer for both liposomal formulations.

### The combined liposomal drug therapy inhibited more effectively the growth of B16.F10 melanoma tumors than each single liposomal drug therapy

To measure the antitumor efficiency of the combined liposomal administration of 5 mg/kg SIM and 14 mg/kg DMXAA in comparison with liposomal monotherapy of either 5 mg/kg SIM or 14 mg/kg DMXAA, drugs were injected intravenously on days 11 and 14 after tumor cell inoculation. The therapeutic agents were also administered as free forms in the same doses and according to the same schedule. Effects of free and liposome-encapsulated drugs on tumor growth were evaluated by measuring the tumor volume at day of sacrification (**Figure 1 A, C, E**) and the area under the tumor growth curve (AUTC) (**Figure 1 B, D, and F**).

**Figure 1:**
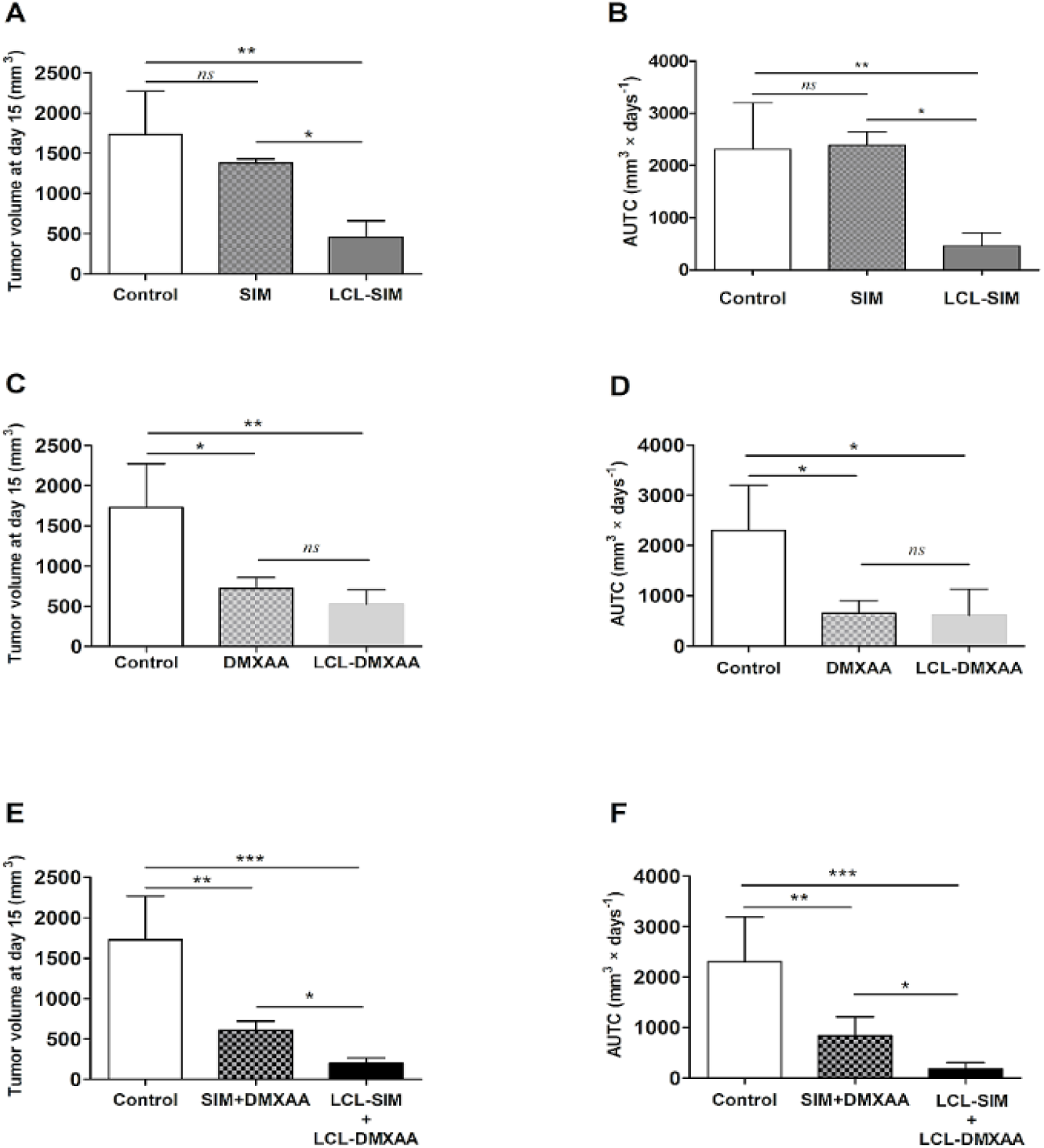
Effects of the combined administration of free and liposome-encapsulated SIM and DMXAA on the growth of *s*.*c*. B16.F10 murine melanoma. Mice received two *i*.*v*. injections of therapeutic agents at day 11 and day 14, after cancer cell inoculation Tumor volumes after different treatments at day 15 (when mice were killed) were presented in panels (**A**), (**C**), and (**E**). AUTCs after various treatments were presented in panels (**B**), (**D**) and (**F**). Control – LCL-treated group; SIM – experimental group treated with 5 mg/kg free SIM; LCL-SIM – experimental group treated with 5 mg/kg SIM as liposome-encapsulated form; DMXAA – experimental group treated with 14 mg/kg free DMXAA; LCL-DMXAA – experimental group treated with 14 mg/kg DMXAA as liposome-encapsulated form; SIM+DMXAA – experimental group treated with 5 mg/kg free SIM and 14 mg/kg free DMXAA; LCL-SIM + LCL-DMXAA – experimental group treated with 5 mg/kg SIM and 14 mg/kg DMXAA as liposome-encapsulated forms. Results were expressed as mean ± SD of tumor volumes of 5 mice. One way ANOVA test with Bonferroni correction for multiple comparisons was performed to analyze the differences between the effects of the treatments on tumor growth (*ns, P*>0.05; *, *P*<0.05; **, *P*<0.01; ***, *P*<0.001).

Our data have shown that LCL-SIM administered alone strongly reduced (by 75-80%, *P*<0.05, **Figure 1 A and B**) melanoma growth compared with administration of free SIM that was totally inefficient in terms of inhibition of tumor growth (P>0.05, **Figure 1 A and B**). This was probably due to the tumor targeting properties of LCL. However, in the case of DMXAA, the encapsulation in LCL did not enhance the antitumor efficacy of the anti-angiogenic drug, since free DMXAA as well as the LCL form exerted similar antitumor activities on tumor growth (60-70% inhibition compared with control, *P*<0.05) (**Figure 1 C and D**). Notably, both combined therapies (free or liposomal) suppressed melanoma growth strongly albeit with much higher degree, by 90-92% inhibition (*P*<0.001) in the case of LCL-SIM+LCL-DMXAA drug therapy and by 60-65% inhibition *(P*<0.01) after administration of free SIM in combination with DMXAA, compared with control tumor growth (**Figure 1 E and F**). However, it is worth to mention that among all therapies tested, the combined liposomal drug therapy exerted the strongest antitumor activity in B16.F10 murine melanoma-bearing mice (**Figure 1 A-F**) (by 25% more tumor growth inhibition compared to free SIM+DMXAA therapy). The effectiveness of the combined liposomal drug therapy on tumor growth might be linked to the tumor targeting properties of the LCL as well as synergistic effect of the combined administration of SIM and DMXAA on melanoma cell proliferation, previously demonstrated by our group [8]. Therefore, the underlying molecular mechanisms of this novel therapeutic approach were further investigated by comparing the effects of single and combined LCL drug administration.

### Combined treatment with LCL-SIM and LCL-DMXAA exerted strong anti-angiogenic effects on B16.F10 murine melanoma *in vivo*

To investigate whether the antitumor activity of the combined liposomal drug therapy can be linked to its action on tumor angiogenesis, a screening for intratumor levels of 24 angiogenic/ inflammatory proteins was performed by protein microarray. In addition, the tumors were analyzed by immunohistochemistry with regard to the expression of CD31, a marker for proliferating endothelial cells [20].

Our results showed (**Figure 2A and Supplementary Table 3**) that the production of almost all angiogenic/inflammatory proteins were inhibited to varying degrees by LCL-SIM (from 9% up to 70%) and by LCL-DMXAA (1-79%) monotherapies compared to Control LCL treated group.

**Figure 2:**
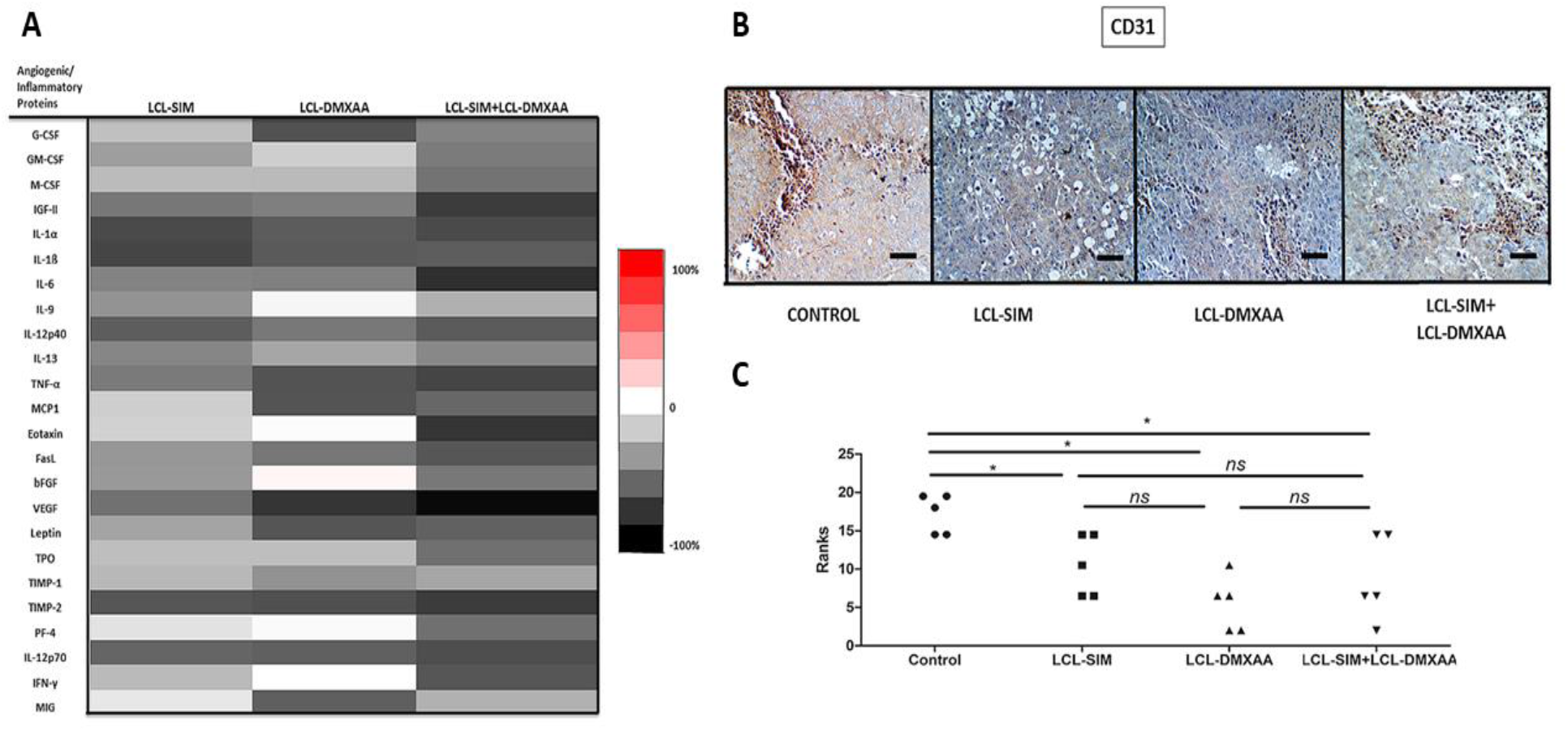
Effects of different treatments on angiogenic and inflammatory protein production and neovascularization in the tumor microenvironment. **(A)**Protein levels after different treatments were compared with the levels of the same proteins in control – LCL-treated group. Data are expressed as average % of reduction (–) of protein levels ranging from 0% (white) to -100% (black) or stimulation (+) of production of proteins ranging from 0% (white) to +100% (red) compared with the levels of the same proteins in control group. **(B)** Immunohistochemical analysis of CD31, a marker for proliferating endothelial cells in *s*.*c* B16.F10 murine melanoma tissue. Positively stained cells for CD31 appear in brown; size bars = 10 μm. **(C)** The scores for immunoreaction intensities were analyzed by using rank-based nonparametric Kruskall-Wallis test with Dunn’s test for multiple comparisons (*ns, P*>0.05; *, *P*<0.05). Control – LCL-treated group; LCL-SIM – experimental group treated with 5 mg/kg SIM as liposome-encapsulated form; LCL-DMXAA – experimental group treated with 14 mg/kg DMXAA as liposome-encapsulated form; LCL-SIM + LCL-DMXAA – experimental group treated with 5 mg/kg SIM and 14 mg/kg DMXAA as liposome-encapsulated forms.

Combined liposomal treatment with SIM and DMXAA exerted the highest suppression (from 30% up to 95%) of the levels of key players involved in tumor angiogenesis and inflammation compared to their control levels. Specifically, compared to their control levels, potent pro-angiogenic/pro-inflammatory proteins such as MCP-1, IL-1α, IL-1β, IL-12p40, TNF-α, Leptin, Fas-L, b-FGF, G-CSF, M-CSF, IL-9, IL-13, GM-CSF were moderately to strongly reduced (by 48-72%), while the levels of IGF-II, a determinant of angiogenic sprouting [21] were reduced by 76% (*P*<0.01) by the combined liposomal therapy. The levels of IL-6, which promotes defective angiogenesis in tumors [22] was reduced by 82% (*P*<0.001) and the levels of eotaxin, a cancer cell invasion promoter [23] were reduced by 79% (*P*<0.01). Notably, VEGF production suffered the most drastic suppression in the combination therapy treated experimental group (>95%, *P*<0.001), in correlation with the strong inhibition of HIF-1α (**Figure 5 A, B**). The average reduction of the proteins by LCL-SIM+LCL-DMXAA treatment was 20% higher compared to LCL-SIM treated group (**Supplementary Table 3 and Figure 2A**). Nevertheless, all anti-angiogenic proteins (TIMP-1, TIMP-2, PF-4, IL-12p70, IFN-γ, MIG) were moderately to strongly inhibited (by 30-76 %) after the combined liposomal treatment with LCL-SIM+LCL-DMXAA.

**Figure 5:**
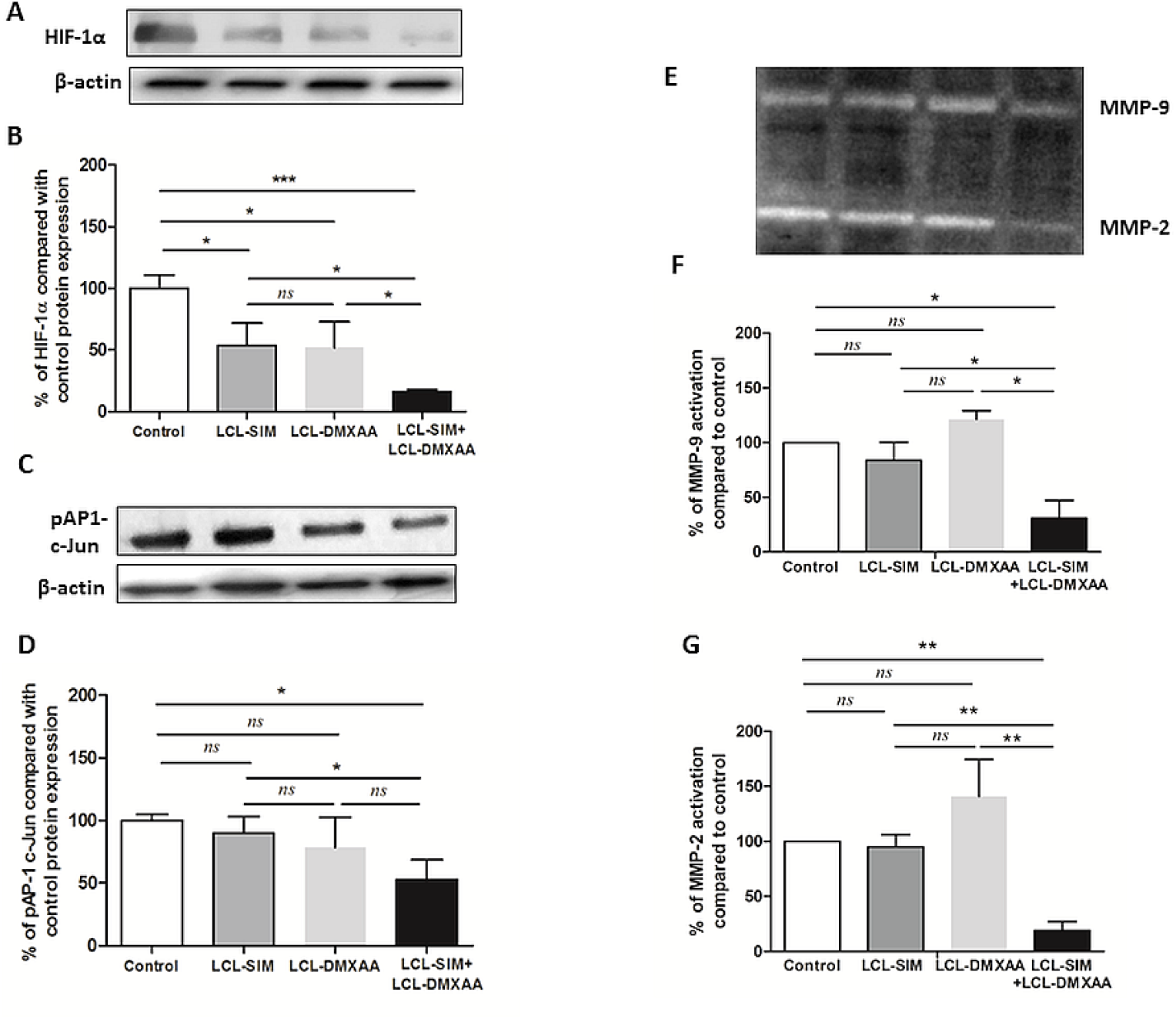
Effects of liposome-encapsulated SIM and DMXAA on intratumoral levels of key invasion and metastasis promoters. **(A), (C)** Western blot analysis showing the effects of different treatments on the intratumor levels of HIF-1α and pAP-1 c-Jun, respectively. β-actin was used as loading control. **(B), (D)** Protein levels of HIF-1α and pAP-1 c-Jun in lysates from treated groups expressed as percentage from Control – LCL-treated group. Data represent the mean ± SD of two independent measurements. One way ANOVA test with Bonferroni correction for multiple comparisons was performed to analyze the differences between the effects of the treatments on apoptotic proteins (*ns*, P>0.05; *, *P*<0.05; ***, *P*<0.001). **(E), (F), (G)** – The effects of different treatments on the activity of microenvironmental matrix metalloproteinases. **(E)** Gelatin zymography analysis of tumor lysates from mice treated with various liposome-encapsulated SIM and DMXAA therapies. Coomassie blue staining highlights gelatinolytic activity corresponding to MMP-9 and MMP-2 (pro-froms and active forms). **(F), (G)** Percentage of MMP-9 and MMP-2 activity in tumor lysates from mice treated with single and combined SIM and DMXAA liposomal therapies compared to control. Data represent the mean ± SD of two independent measurements. One way ANOVA test with Dunnett correction was performed to analyze the differences between the effects of various treatments on MMP-9 and MMP-2 levels compared to control untreated group (*ns, P*>0.05; *, *P*<0.05; **, *P*<0.01). Control – LCL-treated group; LCL-SIM – experimental group treated with 5 mg/kg SIM as liposome-encapsulated form; LCL-DMXAA – experimental group treated with 14 mg/kg DMXAA as liposome-encapsulated form; LCL-SIM + LCL-DMXAA – experimental group treated with 5 mg/kg SIM and 14 mg/kg DMXAA as liposome-encapsulated forms.

Moreover, a significant reduction of CD31 expression in tumors that received treatments based on liposomal drugs compared with its expression in control lysates (**Figure 2B** and **C**, *P*<0.05) was noticed. This finding might suggest a tight connection between inhibition of neovascularization and strong reduction of the production of pro-angiogenic proteins, confirming the strong anti-angiogenic properties of both SIM and DMXAA, already reported by us, *in vitro* [8].

### Co-delivery of liposomal SIM and DMXAA triggers apotosis in B16.F10 melanoma TME

To investigate whether different treatments with liposomal SIM and DMXAA induced apoptosis in melanoma TME, we assessed the relative expression levels of pro-apoptotic Bax and anti-apoptotic Bcl-xL proteins, by western blot. Our results revealed that only the combined treatment with liposomal SIM and DMXAA was able to upregulate Bax protein levels (**Figure 3 A**, **C**, *P* < 0.05) while, all liposomal treatments down-regulated the intratumor production of the anti-apoptotic protein Bcl-xL compared to its level in Control group (**Figure 3 B, D**, *P* < 0.01, *P* < 0.05).

**Figure 3:**
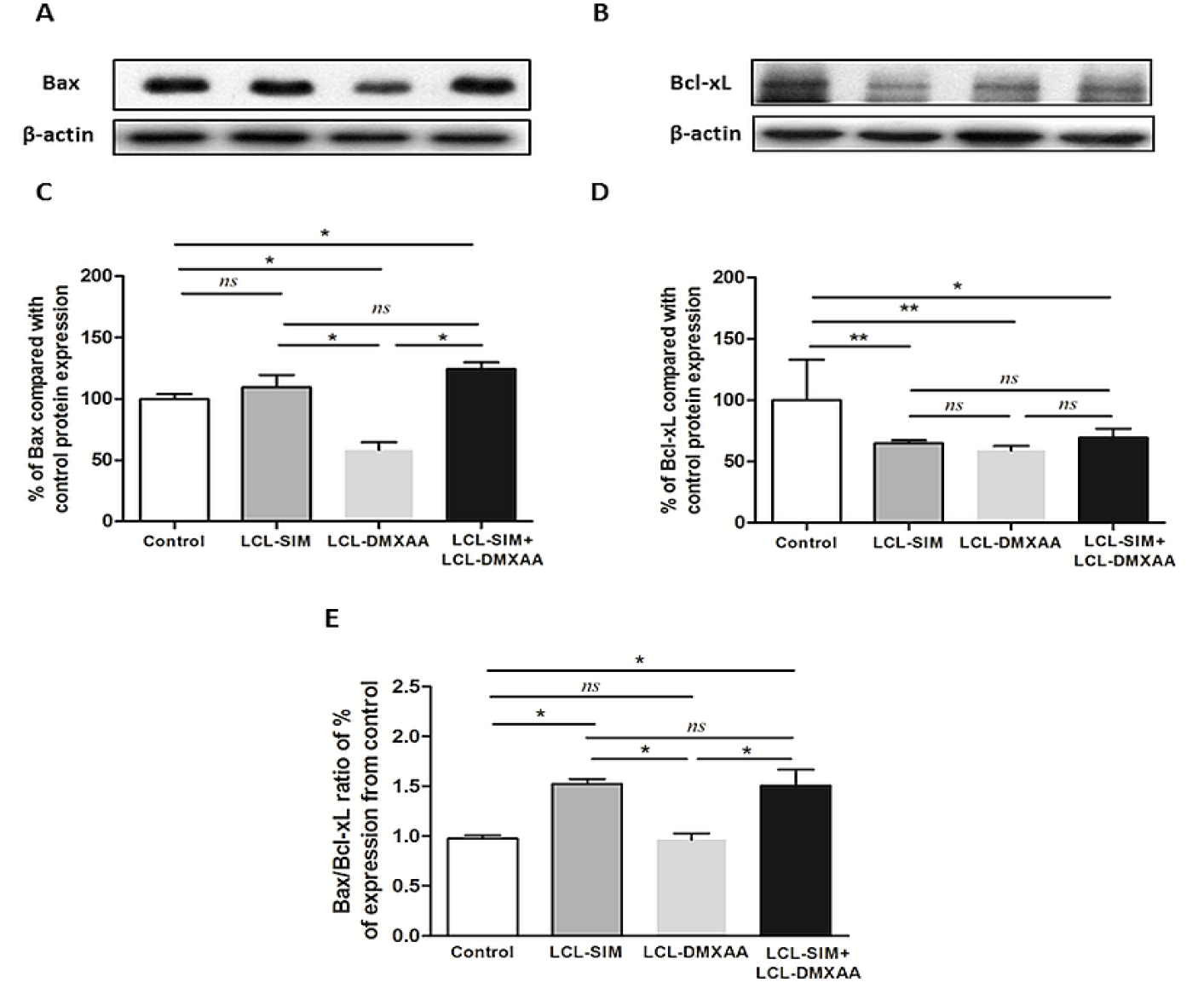
Effects of liposome-encapsulated SIM and DMXAA on intratumoral levels of apoptotic proteins. **(A), (B)**Western blot analysis showing the effects of different treatments on the intratumor levels of Bax and Bcl-xL, respectively. β-actin was used as loading control. **(C), (D)** Protein levels of Bax and Bcl-xL in lysates from treated groups expressed as percentage from control – LCL-treated group. **(E)** The ratio of % expression levels of Bax/Bcl-xL compared to Control. Data represent the mean ± SD of two independent measurements. One way ANOVA test with Bonferroni correction for multiple comparisons was performed to analyze the differences between the effects of the treatments on the apoptotic proteins (*ns, P*>0.05; *, *P*<0.05; **, *P*<0.01). Control – LCL-treated group; LCL-SIM – experimental group treated with 5 mg/kg SIM as liposome-encapsulated form; LCL-DMXAA – experimental group treated with 14 mg/kg DMXAA as liposome-encapsulated form; LCL-SIM + LCL-DMXAA – experimental group treated with 5 mg/kg SIM and 14 mg/kg DMXAA as liposome-encapsulated forms.

Overexpression of Bcl-xL is associated with chemoresistance and metastasis in melanoma and previous studies demonstrated that this protein competes with Bax, negatively influencing the mitochondrial membrane permeabilisation [24,25]. Thus, the Bax/Bcl-xL protein expression ratio was determined and used as a good indicator to estimate the sensitivity of melanoma cells to applied drugs [26]. Our data showed that only LCL-SIM and LCL-SIM+LCL-DMXAA treated tumors showed a 1.5-fold increase in Bax/Bcl-xL production ratio (**Figure 3 E**, *P* < 0.05) indicating that liposomal SIM sensitized melanoma cells to LCL-DMXAA therapy. The histopathological evaluation also revealed some morphologic features of apoptosis induced by all liposomal treatments (**Supplementary Figure 1 A-D**).

### Effects of liposome-encapsulated SIM and DMXAA on intratumor oxidative stress

To link the anti-angiogenic properties of the liposomal therapies with any potential changes in intratumor oxidative stress, the levels of malondialdehyde (MDA), a product of lipid peroxidation, the activity of catalase and TAC were determined in tumor tissue lysates (**Figure 4**). Our results suggested that both single liposomal therapies induced a weak increase in MDA levels (*P*<0.05, **Figure 4A**) whereas the combination liposomal therapy with SIM and DMXAA did not affect MDA levels (*P*>0.05, **Figure 4A**). In addition, a proportional decrease in enzymatic (Catalase) was noticed in tumor lysates from groups treated with LCL-SIM (*P*<0.05), LCL-DMXAA (*P*<0.01) and the liposomal combination therapy (*P*<0.05) (**Figure 4B**). Moreover, except for LCL-SIM treatment which did not affect the TAC (P>0.05), all other treatments with LCL-DMXAA and LCL-DIM+LCL-DMXAA, significantly decreased the levels of Trolox equivalents compared with their level in control tumors (P<0.01 and P<0.05 respectively, **Figure C**).

**Figure 4.**
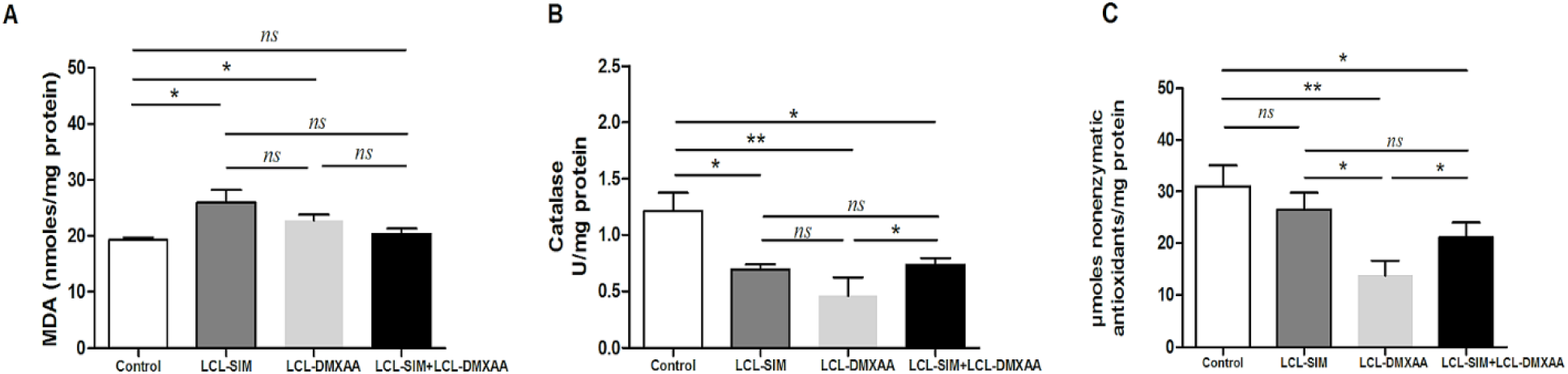
Effects of different liposomal treatments with SIM and DMXAA on tumor oxidative stress parameters. **(A)** MDA concentration expressed as nmoles MDA/mg protein; **(B)** Catalase activity expressed as U/mg protein; **(C)** TAC expressed as μmoles Trolox/mg protein. All parameters were measured in tumor lysates from mice treated with LCL-SIM and LCL-DMXAA as single or combined therapy. Data represent the mean ± SD of duplicate measurements (*ns, P*>0.05; *, *P*<0.05; **, *P*<0.01). Control – LCL-treated group; LCL-SIM – experimental group treated with 5 mg/kg SIM as liposome-encapsulated form; LCL-DMXAA – experimental group treated with 14 mg/kg DMXAA as liposome-encapsulated form; LCL-SIM + LCL-DMXAA – experimental group treated with 5 mg/kg SIM and 14 mg/kg DMXAA as liposome-encapsulated forms.

### Inhibitory effects of the combined liposomal drug therapy with SIM and DMXAA on melanoma invasion and metastasis promoters

The extent of tumor invasiveness and metastasis, after anti-angiogenic therapies depends on the coordinated interaction of numerous proteins and enzymes controlled by several transcription factors such as HIF-1α and AP-1 [27,28]. Thus, we evaluated the impact of the combined liposomal therapy with SIM and DMXAA on intratumor production of metastatic promoters such as HIF-1α and pAP-1 c-Jun, and on the activity of MMP-2 and MMP-9, both of which are activated under hypoxia and degrade the extracellular matrix, facilitating cancer cell dissemination. In line with our previous studies [8], HIF-1α was strongly suppressed by the combined liposomal treatment (80% reduction compared to the protein production in control group) (**Figure 5 A, B**, *P<*0.001), as opposed to single liposomal treatments which only elicited weak inhibitory effects (**Figure 5 A, B**, *P*< 0.05).

Moreover, among all treatments tested, only the combined administration of LCL-SIM and LCL-DMXAA exerted an inhibitory effect on the production of pAP-1 c-Jun (by 47% compared to its control production, **Figure 5 C, D**). Activation of AP-1 is critically linked with Ras-induced oncogenic transformation in melanoma cells and tightly regulates the expression levels of both HIF-1α and MMPs metastatic promoters [29,30]. According to **Figure 5 E-G**, only the combined treatment notably decreased the lytic activity of both MMP-2 (80% inhibition compared to control, *P*<0.01) and MMP-9 (70% inhibition compared to control, *P*<0.05). Together, these results suggest that the reduction of intratumor production of HIF-1α and of pAP-1 c-Jun by combined liposomal therapy with LCL-SIM+LCL-DOX significantly weakened the invasive and metastatic ability of B16.F10 melanoma cells, via strong inhibition of MMPs activity.

### LCL-SIM co-administred with LCL-DMXAA partially “re-educated” TAMs by downregulating the mARN expression of key arginine metabolic enzymes, iNOS and ARG-1

To investigate whether the suppressive effects exerted by combined liposomal therapy on the main TAMs-mediated pro-tumor processes can be linked with the capacity of this therapy to repolarize TAMs towards M1 phenotypes, tumors were evaluated for the expression of iNOS and ARG-1 mARN by RT-qPCR. iNOS and ARG-1 are specific markers for TAMs, a high Arg-1 activity defining the protumor M2 macrophages, while increase iNOS activity is well-recognized as a key biomarker for antitumor M1 macrophages [31]. Our data indicated that LCL-SIM+LCL-DMXAA induced the strongest reduction of mARN expression levels for iNOS (**Figure 6A**, *P*<0.01, 0.41 relative fold change) and ARG-1 (**Figure 6B**, *P*<0.001, 0.17 relative fold change) compared to their mARN expression level in Control group.

**Figure 6:**
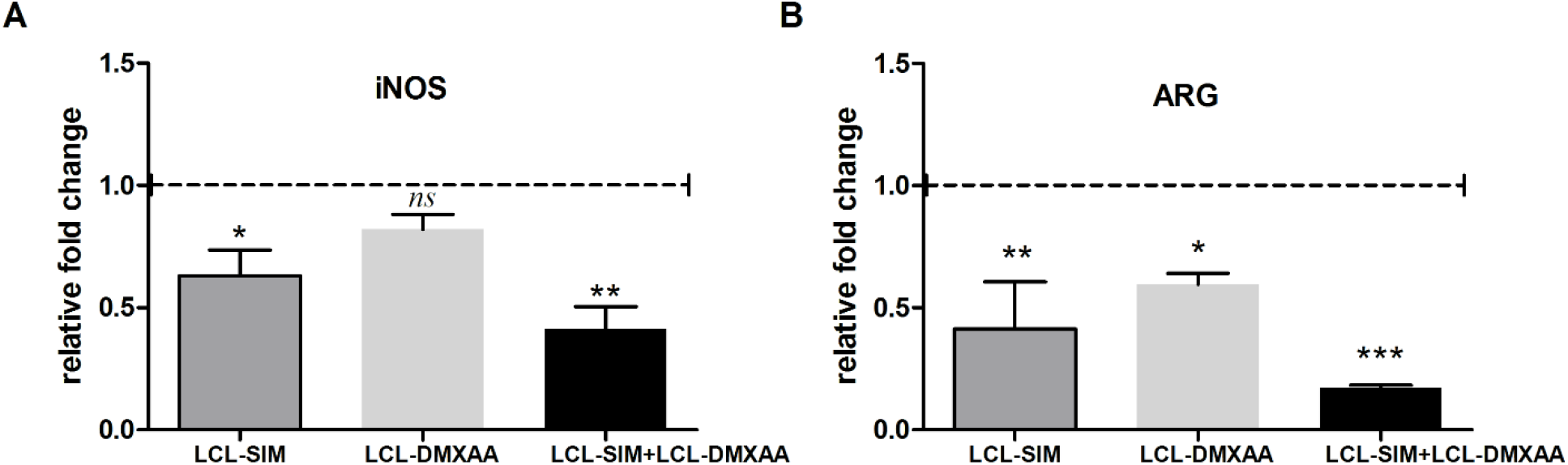
Effects of LCL-SIM and LCL-DMXAA single and combined treatment on the mARN expression levels of enzymes involved in arginine metabolism in TME. **(A), (B)** Effects of liposomal SIM and DMXAA administration on the expression levels of iNOS and ARG-1. mRNA was quantified by RT-qPCR and the results are expressed as fold change based on the Ct calculations. Control – LCL-treated group was used as calibrator. Results were expressed as mean ± SD of three independent measurements. Control – LCL-treated group; LCL-SIM – experimental group treated with 5 mg/kg SIM as liposome-encapsulated form; LCL-DMXAA – experimental group treated with 14 mg/kg DMXAA as liposome-encapsulated form; LCL-SIM + LCL-DMXAA – experimental group treated with 5 mg/kg SIM and 14 mg/kg DMXAA as liposome-encapsulated forms. (*ns, P*>0.05; *, *P*<0.05; **, *P*<0.01; ***, *P*<0.001).

## Discussion

In the present study, we provide a follow-up of our earlier observations that liposomal SIM inhibited melanoma growth *via* concomitant suppressive actions on HIF-1α production in cancer cells and TAMs-mediated oxidative stress [6]. Moreover, these beneficial effects of SIM were recently exploited for the improvement of the anti-angiogenic action of DMXAA on B16.F10 melanoma cells cocultured with TAMs, when these drugs were co-administered [8]. Notably, these data have demonstrated synergistic antitumor action of SIM and DMXAA on melanoma model *via* suppression of the aggressive phenotype of the melanoma cells and TAMs re-education in TME. In the light of these earlier findings, we take the advantage of tumor targeting capacity of LCL to efficiently deliver SIM and DMXAA to B16.F10 melanoma *in vivo* with the final aim of improving the outcome of the anti-angiogenic therapy, which, to our knowledge, has never been described before.

Our study revealed that the administration of LCL-SIM+LCL-DMXAA therapy decelerated almost totally the growth of melanoma *in vivo*, being superior as antitumor efficacy to each liposomal monotherapy tested (**Figure 1A-F**). This beneficial action was tightly connected to the enhancement of the anti-angiogenic action of LCL-DMXAA by LCL-SIM co-administration **(Figure 2A, Supplementary Table 3)**. More specifically, the combined liposomal drug therapy suppressed almost completely the most powerful pro-angiogenic protein-VEGF (with more than 95% compared to its control level) and significantly reduced (p<0.05) the density of blood vessels compared to their density in control tumor tissue (**Figure 2 and Supplementary Table 3**). It is known that altered expression of VEGF, has been found to correlate with melanoma stage and progression [32] and that its targeting appears to offer some therapeutic benefit in melanoma patients, when combined with chemo-or immunotherapy [33]. However, the major drawback of current anti-angiogenic therapies is represented by the hypoxia-induced drug resistance in cancer cells, supported by activation of HIF-1 pathways [34]. As a consequence, a compensatory upregulation of alternative pro-angiogenic molecules occurs in the TME, allowing cancer cells to resist apoptosis and to acquire a more invasive and metastatic behaviour [35]. Therefore, to gain further insight into the LCL-SIM-mediated sensitization of TME to the effects of LCL-DMXAA, we evaluated the impact of this combined therapy on important markers defining TME resistance profile and aggressiveness.

Noteworthy, the combined therapy with LCL-SIM and LCL-DMXAA strongly reduced (by 50-81%) the intratumor production of bFGF, IL-1 α/β, IL-6, TNFα, leptin and of eotaxin, which allow cancer cells to escape VEGF-targeted therapies, promoting tumor growth and metastasis [36,37]. Moreover, it seemed that this combined liposomal therapy did not induce the settlement of cancer cell resistance to anti-angiogenic drugs as treated tumors showed 1.5-fold increase of the Bax/Bcl-xL production ratio (**Figure 3E, P<0.05**) and presented several morphological hallmarks of apoptosis **(Supplementary Figure 1D)**. Other studies previously reported that anti-apoptotic proteins such as Bcl-2 and Bcl-xL are highly expressed in melanoma [38] and a high Bcl-2/Bax or Bcl-xL/Bax ratio correlates with the resilience of cancer cells to undergo apoptosis [26]. Thus, the enhanced intratumor production of pro-apoptotic Bax protein and the strong reduction of intratumor production of IL-6 and Fas-L (**Figure 2A and Supplementary Table 3**), which are of critical importance for cancer cell survival, demonstrate the sensitivity of tumor cells to the combined liposomal therapy and indicate induction of apoptosis [8,12,26,39].

Evidence of accentuated invasiveness of tumor cells following evasive resistance to anti-angiogenic therapy [4], determined us to further assess the effect of our treatments on several established promoters of tumor progression and metastasis (**Figure 5**) such as MMPs. Therefore, the intratumor activity of both MMP-9 and MMP-2, that are involved in facilitating cancer cell dissemination at secondary sites, was investigated by zymography. Although SIM exerts suppressive effects on expression and activation of MMPs [40] and on melanoma cell migration capacity when co-administered with DMXAA [8], our data suggested no significant modulation of MMP-2 and MMP-9 lytic activity by either LCL-SIM or LCL-DMXAA (P>0.05, **Figure 5 E-G**). Notably, when the liposomal drugs were administered concurrently, the activity of both MMP-9 (P<0.05, **Figure 5 E-G**) and MMP-2 (P<0.01, **Figure 5 E-G**) was reduced to a different extent (by 70 % and 80 %, respectively). An explanation for these findings might be given by the similar impact of this treatment on the intratumor production of critical pro-angiogenic/pro-invasive proteins (VEGF, bFGF, IL-1α, IL-1β, IL-6, TNF-α, and eotaxin, **Figure 2A, Supplementary Table 3)** and most importantly, on the production of transcription factors HIF-1α and pAP-1 c-Jun (**Figure 5A-D**). Both HIF-1α and pAP-1 c-Jun proteins are key players in cancer cells resistance to anti-angiogenic therapies, and activate distinct transcriptional programs that converge to coordinate ECM degradation and metastasis, *via* MMPs [29,41]. Thus, the combined liposomal therapy strongly suppressed the intratumor production of HIF-1α (by 84% P<0.001) and of pAP-1 c-Jun (by 47%, P<0.05) proteins compared to their control production (**Figure 5A-D**). Altogether, in spite of the fact that several studies associated the therapeutic inhibition of angiogenesis with increased tumor cell invasiveness [42], our data suggested that by co-delivering LCL-SIM together with LCL-DMXAA, the tumor beneficial association between the activation of HIF-1α/VEGF axis and MMP-2/9 activation [43,44] was blunted.

Our data demonstrated so far that LCL-SIM was able to sensitize the TME to the antitumor actions of LCL-DMXAA. This effect might be associated to the ability of LCL-SIM to partially “re-educate” TAMs [6,8] *via* modulation of arginine metabolism [8]. As shown in **Figure 6B**, when co-administered with LCL-DMXAA, LCL-SIM exerted the strongest reduction of mRNA expression levels of and ARG-1 (P<0.001). These data support our previous published findings [8] regarding the beneficial association of SIM with DMXAA in “re-educating” TAMs towards a M1 phenotype, by reducing ARG-1 expression. Thus, the abolishment of ARG-1 expression in TAMs was associated with the deceleration of the M2 response and inhibition of polyamine synthesis, required for tumor cell proliferation, angiogenesis, cell invasion and metastasis [45] and might mediate the enhancement of cytokine-dependent tumoricidal effects of T cells [46] in tumors. Although our combined liposomal therapy with SIM and DMXAA only partially “re-educated” TAMs towards an antitumor phenotype with regard to iNOS (P<0.01, **Figure 6A**)), inhibiting the expression of mARN coding for both enzymes involved in arginine metabolism might be able to hinder melanoma aggressiveness. Thus, high levels of iNOS-derived NO negatively modulated M1 macrophages differentiation and activation [47] and TAMs-derived iNOS was shown to be instrumental in lymphangiogenesis, an important route for tumor cell dissemination to regional lymph nodes and distant metastasis [48]. Nevertheless, these beneficial suppressive effects on iNOS and ARG-1, might be also be supported by the fact that the combined therapy with liposomal SIM and DMXAA failed to enhance the level of physiological oxidative stress in tumors **(Figure 4A-C)**, therefore inhibiting the settlement of ROS-induced resistance to anti-angiogenic therapy [49,50].

## Conclusions

Taken together, our results showed that the combined liposomal therapy inhibited almost totally the growth of melanoma tumors, due to the enhancement of anti-angiogenic effects of LCL-DMXAA by LCL-SIM and induction of a pro-apoptotic state in the TME. These effects were favored by the partial “re-education” of TAMs towards a M1 antitumor phenotype and maintained via suppression of major invasion and metastasis promoters (HIF-1α, pAP-1 c-Jun, and MMPs) in tumors.

## Author contributions

**Valentin-Florian Rauca, Manuela Banciu:** Conceptualization; **Valentin-Florian Rauca, Manuela Banciu:** Data curation; Formal analysis**; Manuela Banciu:** Funding acquisition**; Valentin-Florian Rauca, Laura Patras, Lavinia Luput, Emilia Licarete, Vlad-Alexandru Toma Alina Porfire, Augustin-Catalin Mot, Alina Sesarman:** Investigation**; Valentin-Florian Rauca, Laura Patras, Lavinia Luput, Emilia Licarete, Vlad-Alexandru Toma Alina Porfire, Augustin-Catalin Mot, Alina Sesarman:** Methodology**; Manuela Banciu:** Project administration**; Manuela Banciu:** Resources**; Valentin-Florian Rauca, Manuela Banciu:** Software**; Manuela Banciu:** Supervision**; Valentin-Florian Rauca, Manuela Banciu, Alina Sesarman:** Validation**; Valentin-Florian Rauca:** Visualization**; Valentin-Florian Rauca, Alina Sesarman:** Writing -original draft**; Manuela Banciu, Elena Rakosy-Tican, Alina Sesarman:** Writing -review & editing.

## Abbreviations

ANOVA: Analysis of variance
ARG-1: arginase-1
AUTCs: Areas under the tumor growth curves
Bax: Bcl-2-associated X protein
Bcl-xL: B-cell lymphoma-extra-large
bFGF: Basic fibroblast growth factor
CD31: cluster of differentiation 31
CHL: cholesterol
DMEM: Dulbecco’s Modified Eagle’s Medium
DMXAA: 5,6-dimethylxanthenone-4-acetic acid
DPPC: 1,2-Dipalmitoyl-sn-glycero-3-phosphocholine; ECM-extracellular matrix
PEG-2000-DSPE: (N-(Carbonyl-methoxypolyethylene-glycol-2000)-1,2-distearoyl-sn-glycero-3-phosphoethanolamine sodium salt)
EPR: enhanced permeability and retention
FasL: Fas ligand
G-CSF: Granulocyte-colony stimulating factor
GM-CSF: Granulocyte-macrophage-colony stimulating factor
HE: Hematoxylin and Eosin
HIF-1α: hypoxia-inducible factor 1α
HPLC: High-Performance Liquid Chromatography
HRP: horseradish peroxidase
iNOS: Inducible nitric oxide synthase
IFN-γ: Interferon γ
IGF-II: Insulin-like growth factor 2
IL-12p40: Interleukin 12 p40
IL-12p70: Interleukin 12 p70
IL-13: Interleukin 13
IL-1α: Interleukin 1α
IL-1β: Interleukin 1β
IL-6: Interleukin 6
IL-9: Interleukin 9
i.v.: Intravenous
LCL: Long circulating liposomes
MCP-1: Monocyte chemoattractant protein-1
M-CSF: Monocyte-colony stimulating factor
MDA: Malondialdehyde
MIG: Monokine induced by IFN-γ
MMPs: Matrix metalloproteinases
MMP-2: matrix metalloprotease-2
MMP-9: matrix metalloprotease-9
pAP-1-c-Jun: Phosphorylated form of c-Jun subunit of AP-1: PBS, Phosphate buffered saline
PF-4: Platelet factor 4
PDI: polydispersity index
ROS: reactive oxygen species
s.c.: Subcutaneous
SD: Standard Deviation
SDS: Sodium dodecyl sulfate
SIM: Simvastatin
TAC: Total antioxidant capacity
TAMs: Tumor-associated macrophages
TBS-T: Tris Buffered Saline with Tween
TIMP-1: Tissue inhibitor of metalloproteinase 1
TIMP-2: Tissue inhibitor of metalloproteinase 2
TME: tumor microenvironment
TNF-α: Tumor necrosis factor α
VDA: vascular disrupting agent
VEGF: Vascular endothelial growth factor

