## Supplemental material for "Remodeling tumor microenvironment by liposomal co-delivery of DMXAA and simvastatin inhibits malignant melanoma progression"

**Supplementary Table 1- Forward and reverse primers used for RT-qPCR**

| Name of genes | | Forward primer (5’-3’) | | Reverse primer(5’-3’) |
| --- | --- | --- | --- | --- |
| Mouse *β-actin* | TCT TTG CAG CTC CTT CGT TGC CGG TCC | | GTC CTT CTG ACC CAT TCC CAC CAT CAC AC | |
| Mouse *ARG-1* | CTC CAA GCC AAA GTC CTT AGA G | | AGG AGC TGT CAT TAG GGA CAT C | |
| Mouse *iNOS* | TTC ACC CAG TTG TGC ATC GAC CTA | | TCC ATG GTC ACC TCC AAC ACA AGA | |

**Supplementary Table 2: Characterization of LCL formulations encapsulating SIM or DMXAA**

| **LCLs** | **Size (nm)** | **PDI** | **Zeta potential (mV)** | **Therapeutic agent concentration** | **Encapsulation**  **efficiency** |
| --- | --- | --- | --- | --- | --- |
| **LCL-SIM** | 135 ± 2.6 | 0.089 | -30 | 1.233 mg/ml | 82.05% |
| **LCL-DMXAA** | 113 ± 3.8 | 0.079 | -41 | 2.367 mg/ml | 39.7% |

**Supplementary Table 3: The effects of LCL-SIM and LCL-DMXAA administered as single as well as combined treatment on TME angiogenic/inflammatory protein production**

| **Angiogenic/**  **inflammatory**  **proteins** | **Percentage of inhibition (-) and stimulation (+) of angiogenic/inflammatory protein production after different treatments compared to control group** | | |
| --- | --- | --- | --- |
|  | **LCL-SIM** | **LCL-DMXAA** | **LCL-SIM+LCL-DMXAA** |
| **G-CSF** | -24.48 ± 3.21 (*ns*) | -67.54 ± 3.23 (*) | -48.55 ± 2.46 (*) |
| **GM-CSF** | -37.85 ± 2.88 (*ns*) | -19.09 ± 4.97 (*ns*) | -51.54 ± 2.76 (*) |
| **M-CSF** | -26.45 ± 1.96 (*ns*) | -26.69 ± 3.59 (*ns*) | -54.67 ± 0.60 (*) |
| **IGF-II** | -53.02 ± 0.21 (*) | -50.08 ± 2.04 (*) | -76.26 ± 1.02 (**) |
| **IL-1α** | -70.92 ± 0.76 (**) | -63.84 ± 1.49 (*) | -70.61 ± 0.25 (**) |
| **IL-1ß** | -72.57 ± 0.37 (*ns*) | -64.40 ± 0.54 (*) | -63.18 ± 1.76 (*) |
| **IL-6** | -48.3891 ± 4.26 (*) | -49.77 ± 2.12 (*) | -81.89 ± 0.95 (***) |
| **IL-9** | -41.97 ± 2.97 (*) | -3.31 ± 1.45 (*ns*) | -30.88 ± 1.97 (*ns*) |
| **IL-12p40** | -63.16 ± 3.06 (*) | -52.78 ± 8.25 (*) | -64.24 ± 0.13 (*) |
| **IL-13** | -47.08 ± 0.66 (*) | -34.86 ± 1.02 (*ns*) | -46.36 ± 1.30 (*) |
| **TNF-α** | -51.77 ± 4.01 (*) | -67.40 ± 13.74 (*) | -72.62 ± 0.62 (**) |
| **MCP-1** | -19.02 ± 0.36 (*ns*) | -66.68 ± 0.91(*) | -59.22 ± 0.57 (*) |
| **Eotaxin** | -17.98 ± 8.20 (*ns*) | -1.08 ± 0.50 (*ns*) | -79.38 ± 5.02 (**) |
| **FasL** | -40.69 ± 1.31 (*) | -53.07 ± 0.66 (*) | -65.52 ± 0.63 (*) |
| **bFGF** | -39.72 ± 1.81 (*ns*) | 3.78 ± 0.82 (*ns*) | -52.63 ± 0.48 (*) |
| **VEGF** | -55.3 ± 3.72 (*) | -79.24 ± 1.62(**) | -95.83 ± 0.14 (***) |
| **Leptin** | -35.57 ± 6.39 (*ns*) | -66.56 ± 1.95 (*) | -61.78 ± 6.35 (*) |
| **Thrombopoietin** | -25.13 ± 0.27 (*ns*) | -25.10 ± 5.84 (*ns*) | -55.40 ± 3.41 (*) |
| **TIMP-1** | -27.51 ± 2.37 (*ns*) | -42.51 ± 4.58 (*) | -34.59 ± 2.59 (*ns*) |
| **TIMP-2** | -65.85 ± 2.06 (*) | -68.50 ± 0.36 (*) | -76.14 ± 0.91 (**) |
| **PF-4** | -11.23 ± 3.04 (*ns*) | -1.77 ± 2.65 (*ns*) | -55.75 ± 0.42 (*) |
| **IL-12p70** | -60.16 ± 2.15 (*) | -62.32 ± 2.4 (*) | -69.02 ± 1.4 (*) |
| **IFN-γ** | -26.82 ± 5.73 (*ns*) | 0.35 ± 0.66 (*ns*) | -65.91±1.33 (*) |
| **MIG** | -9.04 ± 2.13 (*ns*) | -62.72 ± 2.60 (*) | -30.56 ± 28.50 (*ns*) |

*The angiogenic/inflammatory protein levels in tumor lysates after different treatments are compared to control– LCL-treated group levels of the same proteins. The results are expressed as % of the average inhibition (‑) or stimulation (+) ± SD of two independent measurements. The two-way ANOVA Multiple Comparison Test was used to compare overall effects on the production of pro- /antitumor proteins in tumor lysates from all experimental groups ; LCL-SIM – experimental group treated with 5 mg/kg SIM as liposome-encapsulated form; LCL-DMXAA – experimental group treated with 14 mg/kg DMXAA as liposome-encapsulated form; LCL-SIM + LCL-DMXAA – experimental group treated with 5 mg/kg SIM and 14 mg/kg DMXAA as liposome-encapsulated forms (ns, P>0.05; *, P<0.05; **, P<0.01; ***, P<0.001).*

**Supplementary Figure 1**
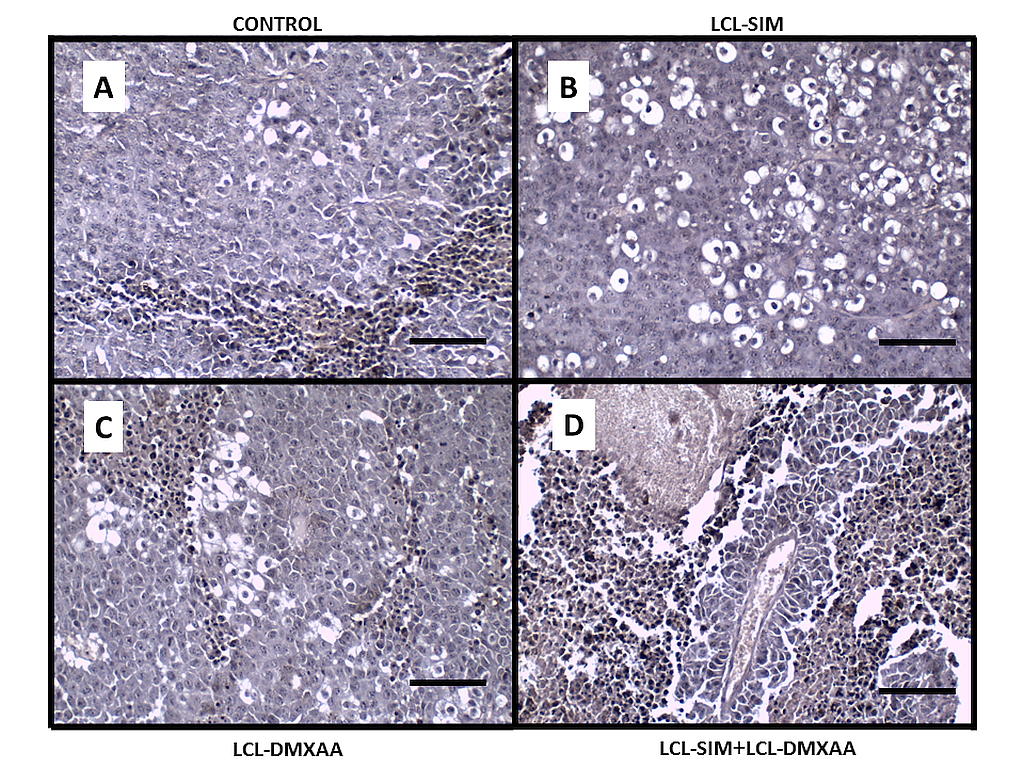

**Supplementary Figure 1: Histopathological evaluation of the effects of different treatments on B16.F10 murine melanoma microenvironment in vivo.** Tissue sections were stained by hematoxylin- eosin method for histological examination; Size bars = 10 μm. **(A)** uncompact histological feature characterized by mitotic nuclei and negligible cytoplasmic vacuolation in tissue sections from Control group; **(B)** intense karyorrhexis and abundant cytoplasmic vacuolation in LCL-SIM-treated tumors; **(C)** moderate inflammatory infiltrate and cytoplasmic vacuolation in LCL-DMXAA treated group**; (D)** intense karyolysis, hyperchorome nuclei and pericapillary infiltration in LCL-SIM+LCL-DMXAA group.
